# Mapping the potential distribution of the principal vector of Crimean-Congo hemorrhagic fever virus *Hyalomma marginatum* in Europe

**DOI:** 10.1101/2022.09.30.510288

**Authors:** Seyma S. Celina, Jiří Černý, Abdallah M. Samy

## Abstract

Crimean-Congo haemorrhagic fever (CCHF) is the most widely distributed tick-borne viral disease in humans. The virus is widely expanded across western China, South Asia, and the Middle East to southeastern Europe and Africa. Its causative agent, Crimean-Congo haemorrhagic fever virus (CCHFV), is among the deadliest human pathogens in Africa and Eurasia. The historical known distribution of the CCHFV vector *Hyalomma marginatum* in Europe included most of the Mediterranean and the Balkan countries, Ukraine, and southern Russia. Further expansion of its potential distribution is possibly occurred in and out of the Mediterranean region. This study updated the map of the principal vector of CCHFV, *H. marginatum*, in the Old World. The model estimated the environmental suitability of *H. marginatum* in the Old World, including Europe. On the continental European scale, the model anticipated a widespread potential distribution, covering southern, western, central, and eastern Europe, as far north as southern parts of Scandinavian countries. The distribution of *H. marginatum* also covered the countries across the central part of Europe where the species is not autochthonous. All models were statistically robust and performed better than random (p < 0.001). Based on the results of the model, climatic conditions could hamper the successful overwintering of *H. marginatum* and their survival as adults in many areas of the region. Regular updates of the models, using updated occurrence, current, and future climatic data are recommended to regularly assess the areas at risk.

**Summary:** *Hyalomma marginatum* is a vector of numerous highly important human and animal pathogens. It is the main vector of Crimean-Congo haemorrhagic fever virus (CCHFV) in Europe. This study updated the potential distribution of *H. marginatum* on a global scale, including Europe, with a particular focus on Central Europe. The model predicted a widespread potential distribution of *H. marginatum* on the continental European scale, anticipating occurrences of *H. marginatum* in southern, western, central, and eastern Europe, as far north as southern parts of Scandinavian countries. In Central Europe, *H. marginatum* populations have not been established in any of the countries yet, but the presence of the species has been reported from all countries in the region. Their potential spread northwards and establishment of permanent populations of *H. marginatum* in the north are therefore of great importance, in particular since the immature stages of *H. marginatum* are frequently found on migratory birds flying northwards to temperate Europe. Our ecological niche model of *H. marginatum* in Central Europe anticipated its distribution in all countries of the region. Our prediction for current global potential distribution of this tick species can help in understanding disease risk areas associated with this vector.

## Introduction

Crimean-Congo haemorrhagic fever (CCHF) is the most widely distributed tick-borne viral disease in humans. The disease is geographically expanded from western China, South Asia, and the Middle East to southeastern Europe and Africa [1]. Crimean-Congo haemorrhagic fever virus (CCHFV) is an emerging arbovirus that causes very dangerous infections in humans with a fatality rate of up to 40% [1–5]. It is also among the deadliest human pathogens in Africa and Eurasia [6].

*Hyalomma marginatum* sensu lato is a complex species including *Hyalomma marginatum* sensu stricto, *Hyalomma rufipes*, and other closely related species [7]. *Hyalomma marginatum* remains the main vector of CCHFV in Europe. *Hyalomma marginatum* has a veterinary and public health importance, particularly if this species can transmit various tick-borne pathogens in humans and animals other than CCHFV such as Spotted Fever rickettsia to humans, *Anaplasma* species to animals, *Babesia caballi* and *Theileria equi* (piroplasmosis) to horses, and *Theileria annulata* (tropical theileriosis) to bovines [8–11]. *Hyalomma marginatum* is one of the tick species whose distribution and expansion are closely monitored by the European Center for Disease Prevention and Control (ECDC) owing to its major medical importance [12].

*Hyalomma marginatum* is well adapted to a wide range of abiotic conditions but it prefers rather arid localities with high summer temperatures [13]. *Hyalomma marginatum* is a two-host tick where the species can feed on a wide range of different host species. Immature developmental stages feed on the same individual animal (e.g., a small mammal like hares, hedgehogs, and rodents) or a ground-dwelling bird, while adults prefer larger animals (e.g., cattle and horses) or occasionally humans [13]. Large domestic mammals play an important role in *H. marginatum* biology and transmission of *Hyalomma*-borne pathogens by supporting a high tick load and bringing *H. marginatum* into proximity to agricultural workers. Additionally, livestock can directly expose humans to *Hyalomma*-borne pathogens through infected blood or crushing of engorged ticks on the animals during slaughtering [14–16]. Migratory birds may have a potential role in the introduction of infected *H. marginatum* into new geographic regions by carrying immature ticks during feeding [13].

Recently, *Hyalomma marginatum* was identified in Ukraine and southern Russia [7]. The wide dispersion of *H. marginatum* reflects their tolerance to diverse environments, including savannah, steppe, and forest steppe, and the ability of their aggressively questing larvae and nymphs to feed on a variety of hosts [13]. *Hyalomma marginatum* ticks prefer the Mediterranean climate of North Africa and southern Europe with low to moderate levels of humidity and a long dry season during the summer months.

*Hyalomma marginatum* was first detected in Central Europe in southern Germany in 2007 [17]. *Hyalomma marginatum* were recorded in other countries of Central Europe like Hungary in 2009 [18], Slovakia in 2008-2009 [19], Austria in 2018 [20], and Czech Republic in 2018-2019 [21]. Permanent populations of *H. marginatum* are generally limited to the warmer areas of the Mediterranean basin in Europe but the occurrence of their permanent populations in Central Europe is probably occurred due to ongoing climatic changes. The establishment of permanent populations and northern spread of *H. marginatum* are anticipated due to passive transportation of immature stages by migratory birds flying to temperate Europe [22,23]. During the spring migration of migratory birds from the south to the north, *H. marginatum* is possibly introduced to Central Europe [24–27]. The spread of these ticks into new geographic locations may possibly cause the emergence of CCHFV in these novel invaded areas. The presence of the virus, its vectors, reservoirs, and amplifying hosts are necessary for the emergence of CCHF in the presence of suitable environmental conditions [28]. Climate change and international animal trades have also significant impacts on modifying the distributional potential of *H. marginatum* and allowing the emergence of CCHF into new geographic regions such as Central Europe.

Here, we aimed to assess the potential distribution of *H. marginatum*, the principal vector of CCHFV, in Europe, with a particular focus on Central Europe by using ecological niche modeling approach. Thus, it derived the detailed pictures to assess its potential to invade new areas in Central Europe.

## Materials and Methods

### Occurrence Records

The occurrence records of *H. marginatum* were obtained from the Global Biodiversity Information Facility (GBIF; www.gbif.org). These datasets were subjected to several data cleaning steps by removing duplicate records to reduce possible biases in estimating ecological niche models [29,30]. Occurrence records were filtered based on a distance filter of ≤2.5’ (≈ 5 km) using SDMtoolbox 2.4 [31] in ArcGIS 10.7.1 (Environmental Systems Research Institute (ESRI), Redlands, CA). The final thinned occurrence dataset of GBIF was divided randomly into two portions using Hawth’s Tools [32] available in ArcGIS 10.7.1: 75% for model calibration, and 25% for internal evaluation of model predictions.

### Accessible Area (*“M”*)

The accessible area ***‘’M”*** is an important element in the biotic, abiotic, and movement (***BAM***) diagram [33] and defines the key parameters in constructing an ecological niche model for the species in question [34]. Accessible area ***“M”*** indicates the areas that the species explored and had access to over relevant periods of the species’ history [34]. The delimitation of *H. marginatum* accessible area *‘’M”* was estimated using *grinnell* package in R [35]. This method simulates dispersal and accessibility based on niche estimations [35].

### Covariate Variables

Several sets of environmental variables were used as independent variables in our model to characterize environmental variations across the calibration and projection areas. These variables were defined as potential drivers limiting the distributional potential of the vector *H. marginatum* [36]. These variables included satellite data from WorldGrids [37], daytime and nighttime land surface temperature (LST), enhanced vegetation index (EVI), and Topographic Wetness Index (TWI). EVI data was included due to its significant role in shaping the ecological niches of tick vectors, particularly immature stages which live on vegetation [36]. EVI is also considered an important factor in reflecting soil moisture’s availability for larvae and nymphs [38,39]. Humidity and aridity are also important factors, considering their major roles in completing the life cycle of *H. marginatum* ticks. Effects of abiotic factors such as temperature and humidity on the distribution of *H. marginatum* have been previously described and used in modeling diverse tick vector species [40,41]. To account for these factors, we used data summarizing climate variables from CHELSA database (http://chelsa-climate.org/) in 30 arc sec spatial resolution (≈ 1 km). CHELSA data includes 19 bioclimatic variables originally derived from monthly temperature and rainfall values collected from weather stations from 1981-2010 [42]. Bioclimatic variables 8-9 and 18-19 were excluded from the analysis to avoid problems deriving from odd spatial artifacts [29].

Anthropogenic data consisted of data on human population density from 2015-2020, nighttime lights, and accessibility via transportation that might play role in the distribution of *H. marginatum* ticks. Population density grids were obtained from the Gridded Population of the World, version 4 (GPWv4) available at http://beta.sedac.ciesin.columbia.edu/data/collection/gpw-v4 [43]. Nighttime satellite imagery was obtained from the NOAA-Defense Meteorological Satellite Program (http://ngdc.noaa.gov/eog/dmsp/downloadV4composites.html) and used as a proxy for real poverty [44,45]. Accessibility was summarized in terms of travel time by land or sea [46], as the connectivity between population sites, is an important variable in estimating the potential distributions of disease vectors and emerging diseases [30,47]; this layer was developed by the European Commission and World Bank (http://forobs.jrc.ec.europa.eu/products/gam/download.php). Final sets of variables were as follows: Set 1 (15 bioclimatic variables from CHELSA); Set 2 (12 variables selected based on jackknife test in MaxEnt); Set 3 (a combination of 15 bioclimatic variables and satellite data summarizing daytime and nighttime LST, EVI, TWI, population density, nighttime satellite imagery, and accessibility); and Set 4 (only satellite data summarizing daytime and nighttime LST, EVI, TWI, population density, nighttime satellite imagery, and accessibility) (**Table 1**).

**Table 1.** Settings and variables used for the construction of the ecological niche modeling for *H. marginatum*. Settings and model performance under optimal parameters using sets of environmental predictors (SEP), statistically significant models (SSM), best candidate models (BCM), regularization multiplier (RM), features classes (FC), mean Area Under the Curve ratio (AUC.r), partial Receiver Operating Characteristic (p.ROC), omission rate 5% (O.rate 5%), Akaike information criterion corrected (AICc), delta Akaike information criterion corrected (ΔAICc), Akaike information criterion corrected weight (AICc.W), number of parameters (#; summarizes the combination of environmental variables, multiple regularizations, and features other than 0 that provide information for the construction of the model based on lambdas), and candidate sets of environmental variables tested during calibration of *H. marginatum* model.

| SEP | SSM | BCM | RM | FC | AUC.r | p.ROC | O.rate 5% | AICc | ΔAICc | AICc.W | # |
| --- | --- | --- | --- | --- | --- | --- | --- | --- | --- | --- | --- |
| Set4 | 1846 | 1 | 3.0 | lph | 1.21 | 0.00 | 0.05 | 1960.75 | 0.00 | 1.00 | 15 |
| Candidate sets of environmental variables of *H. marginatum* model |  |  |  |  |  |  |  |  |  |  |  |
| Set1 |  |  | Set2 |  |  | Set3 |  |  | Set4 |  |  |
| Bio 1, Bio 2, Bio 3, Bio 4, Bio 5, Bio 6, Bio 7, Bio 10, Bio 11, Bio 12, Bio 13, Bio 14, Bio 15, Bio 16, and Bio 17 |  |  | Bio 1, Bio 2, Bio 3, Bio 4, Bio 5, Bio 6, Bio 7, Bio 10, Bio 11, Bio 14, Bio 15, Bio 17 |  |  | Bio 1, Bio 2, Bio 3, Bio 4, Bio 5, Bio 6, Bio 7, Bio 10, Bio 11, Bio 12, Bio 13, Bio 14, Bio 15, Bio 16, Bio 17, Daytime and Nighttime LST, EVI, TWI, Population density, Nighttime satellite imagery, and Accessibility |  |  | Daytime and Nighttime LST, EVI, TWI, Population density, Nighttime satellite imagery, and Accessibility |  |  |

Bio 1: Annual Mean Temperature; Bio 2: Mean Diurnal Range; Bio 3: Isothermality; Bio 4: Temperature Seasonality; Bio 5: Maximum Temperature of Warmest Month; Bio 6: Minimum Temperature of Coldest Month; Bio 7: Temperature Annual Range; Bio10: Mean Temperature of Warmest Quarter; Bio11: Mean Temperature of Coldest Quarter; Bio12: Annual Precipitation; Bio13: Precipitation of Wettest Month; Bio14: Precipitation of Driest Month; Bio15: Precipitation Seasonality; Bio16: Precipitation of Wettest Quarter; Bio17: Precipitation of Driest Quarter; LST: Land Surface Temperature ; EVI: Enhanced Vegetation Index; and TWI: Topographic Wetness Index.

### Ecological Niche Modeling of *H. marginatum*

We constructed ENM using the maximum entropy algorithm implemented in MaxEnt version 3.4.1 via the *kuenm* R package [48]. 1972 candidate models, with parameters reflecting all combinations of 17 regularization multiplier settings (0.1-1 with intervals of 0.1, 2-6 with intervals of 1, and 8 and 10), 29 possible combinations of 5 feature classes (linear = l, quadratic = q, product = p, threshold = t, and hinge = h), and 4 distinct sets of environmental and socioeconomic variables were created. The best candidate model was selected based on three different criteria: 1) significance, 2) performance, and 3) the Akaike information criteria (AIC): AICc, delta AICc, and AICc weights. Performance was measured using omission rate which is a threshold that considers an estimate of the likely amount of error among occurrence data and thus removes 5% of occurrences with the lowest suitability values (E = 5%) [49]. Models were selected with delta AICc≤2 from those that were statistically significant and had omission rates below 5%. We followed the criteria from the original *kuenm* study [48] to select the final model, and evaluating the model. We created the final model of *H. marginatum* using 10 replicates by bootstrap, with logistic, product, and hinge outputs. These models were finally transferred from the accessible area ***“M”*** to the projection area ***“G”***.

Model performance was evaluated based on statistical significance (partial ROC), omission rates (OR), and the Akaike information criterion (AICc). Partial ROC and omission rates were evaluated based on models created with training occurrences, whereas AICc values were calculated for models created with the full set of occurrences [50].

### Extrapolation Risk of *H. marginatum*

This analysis identified the areas with extrapolation risk based on a mobility-oriented parity (MOP) approach to compare the environmental breadth of predictors within ***“M”*** (10% reference points sampled) with that in the projection area. The MOP analysis was performed using the MOP function [51] available in the *kuenm* R [48]. The risk of extrapolation analysis calculates multivariate environmental distances between projection area ***“G”*** and the nearest portion of the calibration region to identify areas that have a condition of strict or combinational extrapolation.

### Independent Evaluation

We used a set of additional independent records. These records were retrieved from the previous study [36] for additional evaluation of the model performance to assess its ability to anticipate risk in unsampled areas. We cleaned the dataset to keep only unique records and double-checked the records to remove any of source of overlap with the data used in the model training. This analysis used a one-tailed cumulative binomial probability test to assess the probability of obtaining the observed level of correct predictions by chance alone given the background expectation of correct predictions determined by the proportional coverage of the study area by regions of predicted suitability.

## Results

The final dataset included 716 occurrence points (95 points for model calibration and 621 points for independent evaluation), after removing duplicated occurrences and redundant occurrence records (**Figure 1**).

**Figure 1.**
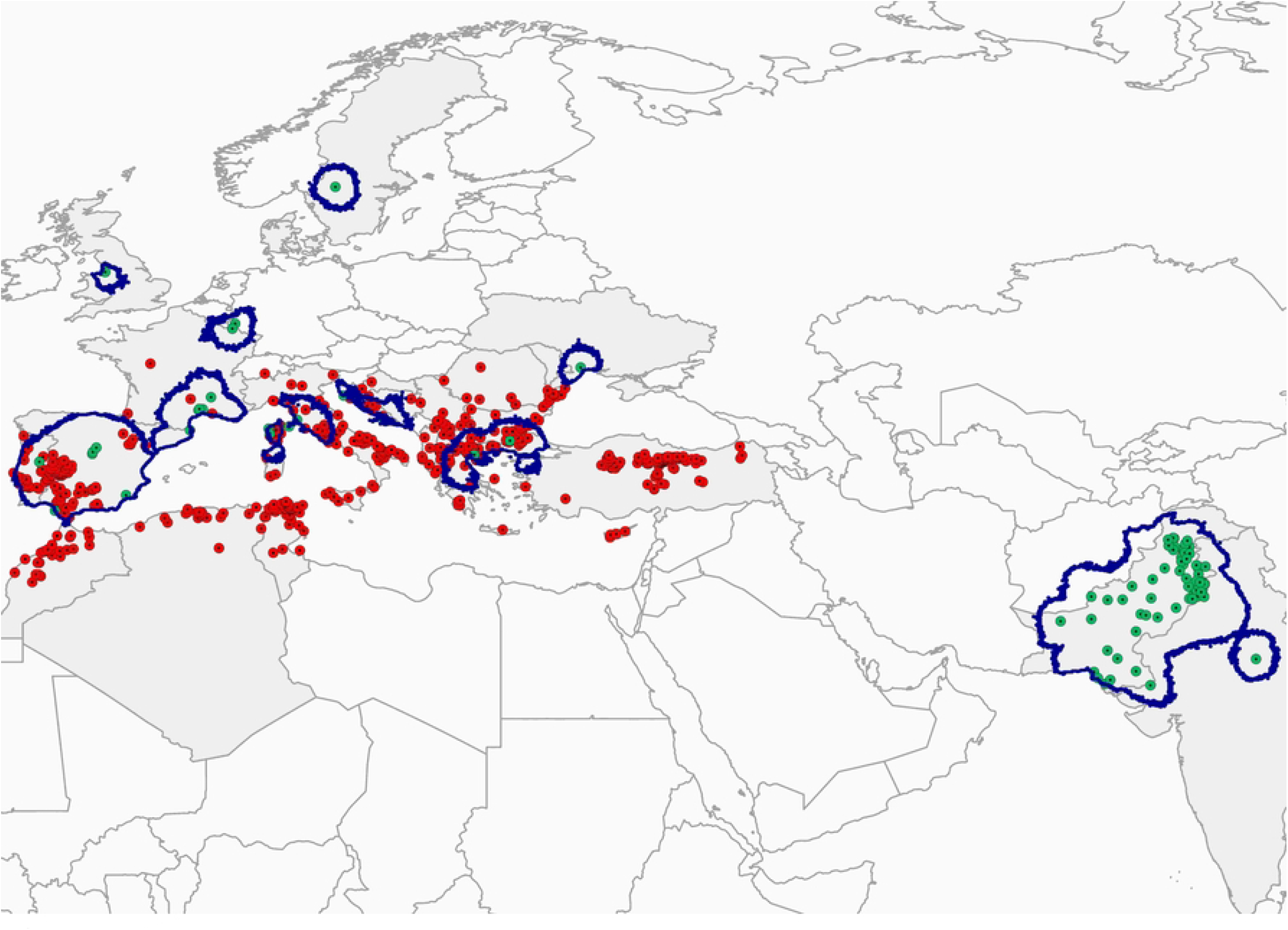
*H. marginatum* occurrence records used in model calibration and final model evaluation across Europe, North Africa, Western and South-Central Asia. The green dotted circles represent the occurrence records of *H. marginatum* collected from GBIF used for model calibration across the ***‘’M”*** area. The red dotted circles represent the retrieved occurrence records of *H. marginatum* from the literature used for the final model evaluation. The gray background depicts the countries where the *H. marginatum* occurrences used in the modeling were recorded. Blue boundaries represent the accessible areas (‘’***M***”) where the *H. marginatum* model was calibrated.

1846 models were statistically significant out of 1972 candidate models. Finally, only one model met the three selection criteria and was identified as the best candidate model based on its performance (**Table 1**). The best model used set 4 of satellite data summarizing Daytime and nighttime land surface temperature, EVI, TWI, Population density, Nighttime satellite imagery, and Accessibility.

In Africa, northern parts of North Africa (Morocco, Tunisia, and Egypt), West and East Africa, central parts of Central Africa, and eastern parts of Southern Africa presented high to very high suitable conditions for *H. marginatum* occurrence (**Figure 2**). In Asia, the model depicted medium to high suitability in expanded large areas of the continent (including India, Pakistan, Bangladesh, and parts of China) (**Figure 2**).

**Figure 2.**
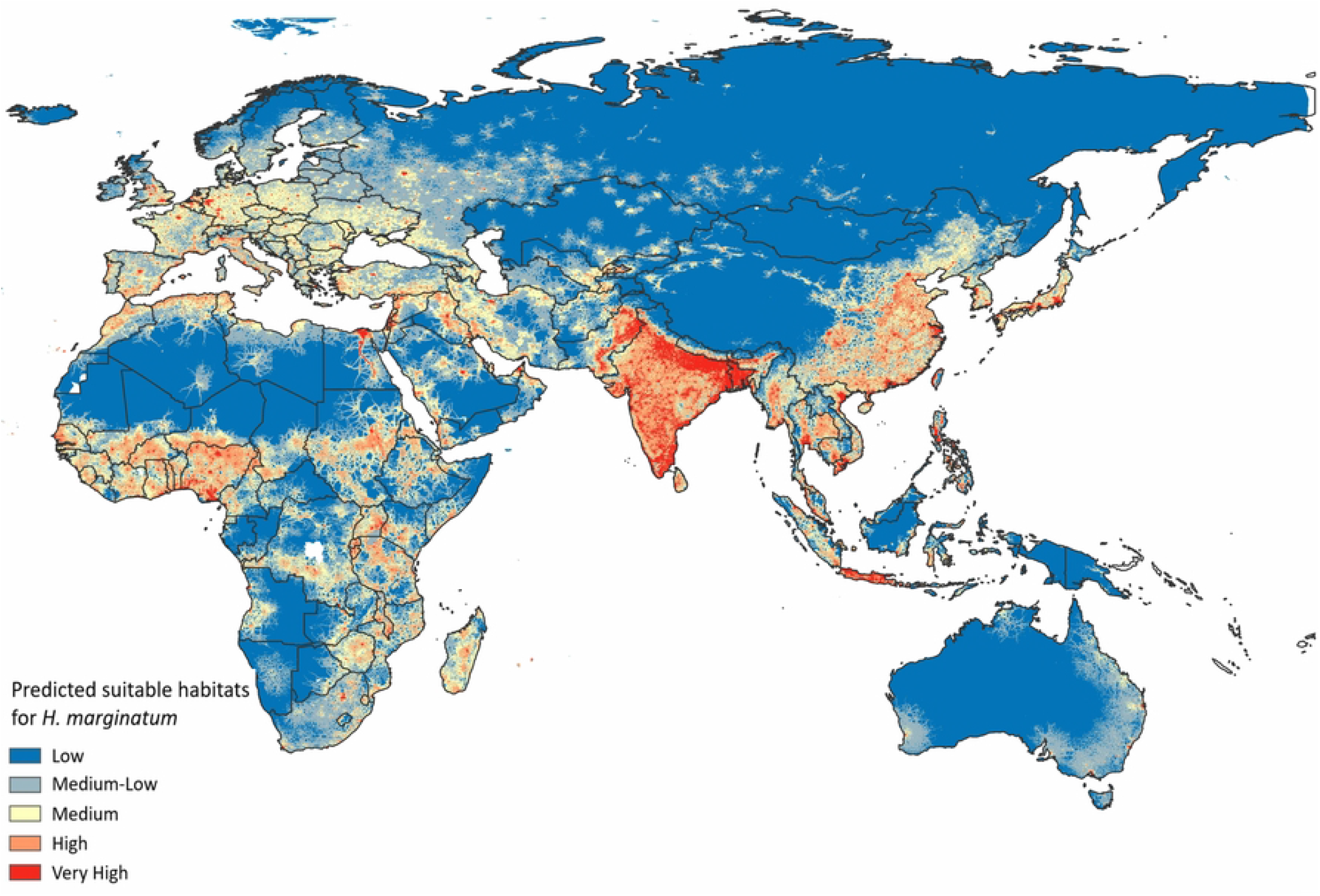
Predicted potential geographic distribution of Crimean-Congo haemorrhagic fever vector *H. marginatum* on a global scale. Red colors indicate highest habitat suitability and blue lowest suitability.

The model anticipated a widespread potential distribution on the continental European scale (**Figure 3**). Thus, the model anticipated occurrences of *H. marginatum* in southern, western, central, and eastern Europe, as far north as southern parts of Scandinavian countries (**Figure 3A**). A high probability of suitable conditions for *H. marginatum* in southern Europe, especially in the Mediterranean parts of Spain, Portugal, Italy, and Greece and along the Adriatic shore was presented. In the Balkan Peninsula, the southeastern part of the continent, wide areas of all countries are identified as highly suitable areas for *H. marginatum*. In western Europe, large parts of northern and southern France and the Benelux states are predicted to be highly suitable for the distribution of the tick species. Even though all Scandinavian countries broadly present low suitability for *H. marginatum*, the model predicted medium to high suitability in southern regions of all most north European countries (Sweden, Norway, Denmark, and Finland except Iceland), as well as high suitability in scattered areas in southern parts of north European countries. Large parts of the northeastern region of Europe demonstrated low and low-medium suitability for *H. marginatum* distribution, whereas only a few scattered points indicated medium to high suitability in Baltic countries (Estonia, Latvia, and Lithuania). In northwestern Europe, the model depicted medium suitability across all areas in the UK, and very high suitability in Southeast and Southwest of England, East Midlands, West Midlands, Yorkshire, London, southern parts of Northwest, and central parts of Northeast of England. In Scotland, Ireland, and Northern Ireland, low and low-medium suitability for *H. marginatum* distribution predominates. Eastern Europe is identified as a highly suitable area, covering wide medium and very high suitability across Romania, Ukraine, Moldova, and the central part of Moldova.

**Figure 3.**
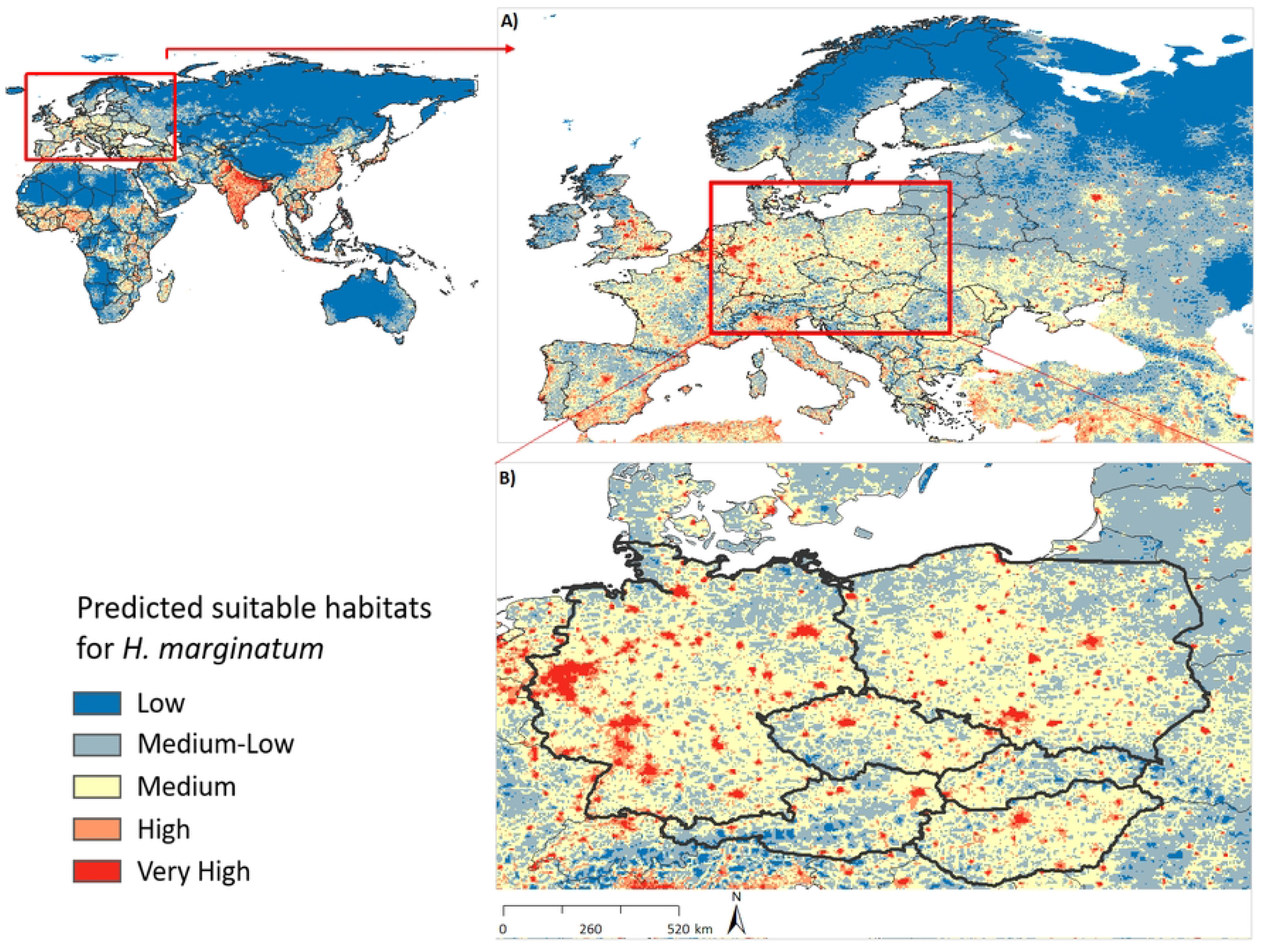
Predicted potential distribution of Crimean-Congo haemorrhagic fever vector *H. marginatum* on a global scale (left top), and close-ups of Europe (A) and Central Europe (B), to provide additional detail to predictions in the region. Red areas indicate modeled highest suitable conditions, and blue areas are lowest suitable conditions.

The ENM of *H. marginatum* in Central Europe anticipated its distribution in all countries of Central Europe (**Figure 3B**). The model predicted broader environmental suitability in Germany, Poland, Hungary, and Czech Republic, followed by Austria and Slovakia. Medium to high suitability is depicted throughout Germany, with the highest suitability across North-Rhine Westphalia, Hesse, Saarland, Baden-Württemberg, Bremen, Berlin, Hamburg, and many scattered points in Bavaria, Thuringia, Saxony, and Lower Saxony. The same is true for large parts of Poland and Hungary, particularly the entire Budapest was identified as a very high suitable habitat for *H. marginatum* occurrence. There are also high-risk areas occurred in eastern Austria and adjoining areas, as well as western and northern parts of Slovakia. While large areas of Czech Republic have medium suitability, very high suitability of *H. marginatum* also occurred in several scattered areas.

Final model was evaluated by using independent occurrence data [36]. The independent set of *H. marginatum* occurrence records were related to the *H. marginatum* model prediction. The model successfully predicted 588 of 621 (94.68%) of these independent data (**Figure 4**).

**Figure 4.**
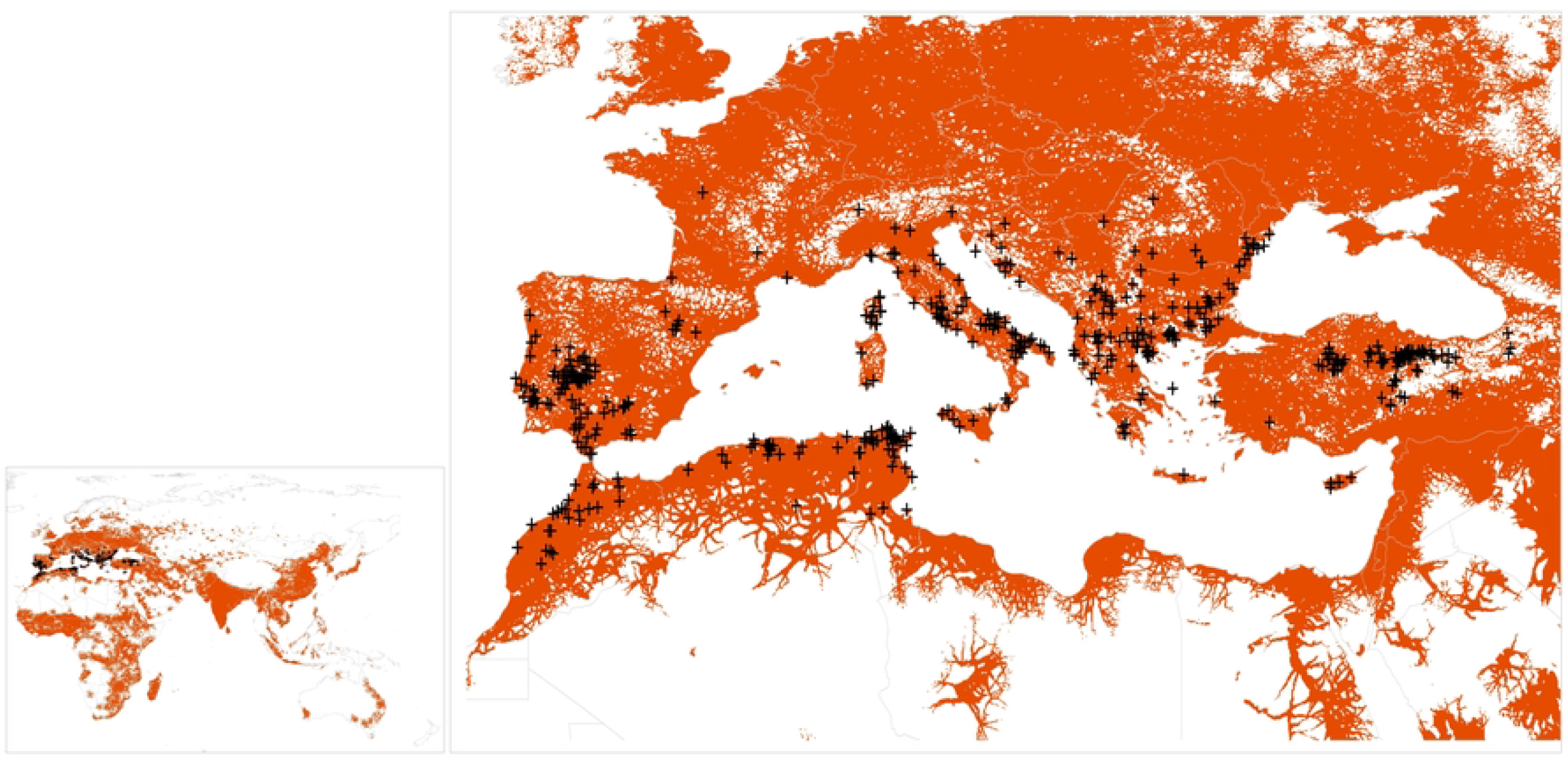
Relationship of *H. marginatum* ecological niche modeling prediction to the distribution of the independent set of *H. marginatum* occurrence records. Black plus signs represent the retrieved occurrence records of *H. marginatum* from the literature used as an independent subset of occurrence data for the final model evaluation; orange shading shows areas predicted by models as suitable for *H. marginatum* distribution.

The MOP results indicated high levels of environmental similarities in all areas under the question, except Southwest China (e.g., northern parts of Tibet province), Northwest China (e.g., southern parts of Xinjiang province), and some areas in East Africa (e.g., Ethiopia, Kenya, Somalia, Sudan) where strict extrapolation occurred (**Figure 5**). Therefore, predictions on these areas should be taken with caution as they were consistently detected as areas with high extrapolation risk.

**Figure 5.**
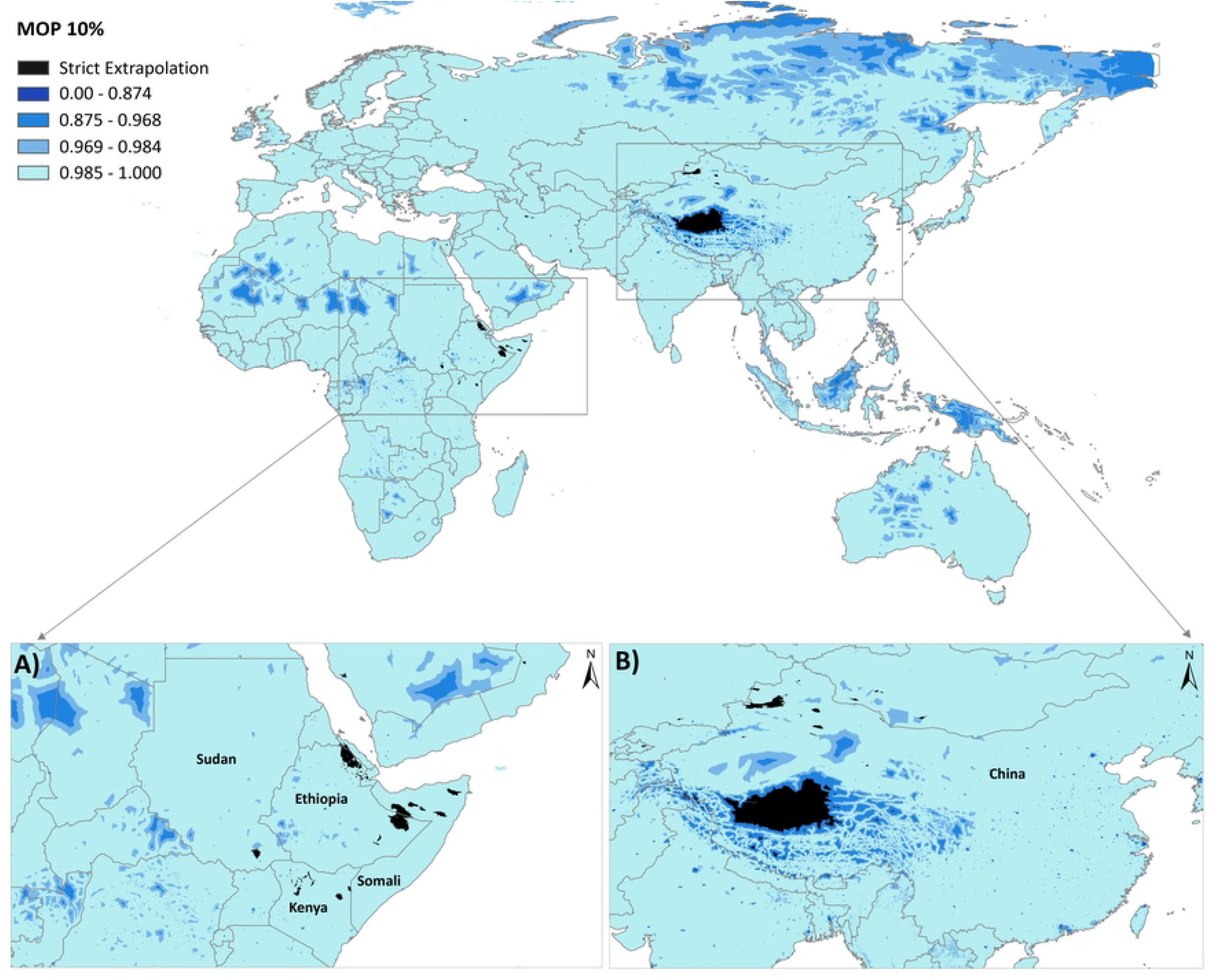
Mobility-oriented parity (MOP) 10% extrapolation risk analysis for the ecological niche model of *H. marginatum* from the calibration area (*“M”*) to a projection area (*“G”*) (top), and close-ups of East Africa (A) and Eastern Asia (B), to provide additional detail to strict extrapolations occurred in the areas. The MOP analysis indicated that areas with the most dissimilar variables conditions (i.e., where one or more covariate variables are outside the range present in the training data) were found beyond the potential distributional areas predicted by the model in the ***“G”*** area. Areas with the most dissimilar variables conditions display strict extrapolative areas and are represented by zero value. Other values represent levels of similarity between the calibration area and the ***“G”*** transfer area. The MOP raster output was reclassified into five categories; the first category represented a strict extrapolation (i.e., zero value), and the fifth category represented the highest environmental similarities between calibration and projection areas.

## Discussion

Ecological niche modeling has been widely implemented to successfully predict the distribution of numerous vector-borne diseases [30,52–60]. These modeling studies leads to more effective implementation of vector surveillance and control programs.

Here, the study provided detailed maps to identify the potential distribution of *H. marginatum*, principal vector of CCHFV, in the Old World, with a particular focus on Europe. The study also provided additional estimates of model uncertainty under present environmental and socioeconomic conditions. These conditions identified several key factors contributing to the spread of ticks and tick-borne diseases such as climate change, vegetation, human population, poverty, and increased animal and human movement via accessibility. The introduction of *Hyalomma* species into new geographical areas is probably due to the presence of infested hosts resulting from human activity, movement of farm and wild mammals, and migrating birds [8,61].

*H. marginatum* is a non-nidicolous tick vector that is actively seeking hosts using an active locomotory hunting strategy, which contrasts with the *Ixodes* ticks which passively wait in the vegetation for a vertebrate host to come accidentally into contact with them (ambush strategy) [62]. After spotting a host by sensing certain signals such as vibration, visual objects, carbon dioxide, ammonia, or body heat, *H. marginatum* can run rapidly several meters across the ground to attack the host. Therefore, they are known as “hunter ticks”. *H. marginatum* has close associations with a wide range of vertebrate hosts including small and large mammals, birds, and reptiles [13]. Based on its ability of using numerous vertebrate hosts, no host parameters were included in the model since good quality occurrence data for numerous host species is difficult to acquire and it is unclear how best to incorporate them [41]. *H. marginatum* is increasingly found outside of its known range in many parts of Europe, possibly due to changes in climate and changes in animal and human movements. Thus, anthropogenic data is included in the model to provide a more reliable prediction of this vector species.

*H. marginatum* is a vector of numerous highly important human and animal pathogens including CCHFV. The distribution of CCHFV is related to the distributional potential of its vector species. Climate change impacts, transportation of immature ticks through international animal trade, and migratory birds have a crucial role in geographic expansion of CCHFV.

Historical known distribution of *H. marginatum* in Europe includes most of the Mediterranean countries, Ukraine and southern Russia. Further expansion of its potential distribution is possibly occurred in and out of the Mediterranean region. In Europe, *H. marginatum* is endemic mostly in Mediterranean but isolated populations exists also in Transalpine Europe. Our study showed that large areas are highly suitable for *H. marginatum* occurrence across entire Europe, particularly the Mediterranean parts of Spain, Portugal, Italy, Greece and along the Adriatic shore, southern and northern France, large areas in Netherlands, Belgium, Germany, Balkan countries, Ukraine and Crimea, scattered areas across countries of Central Europe, and extending well into United Kingdom (**Figure 2**, **Figure 3**). A previous study anticipated high environmental suitability of *H. marginatum* across Southern, Southeastern and Central Europe, North Africa near the northern coast of Tunisia, Morocco, and Algeria, across Arabian Peninsula and south-central Asia, and China [63]. In the current study, we updated the ENM of *H. marginatum* to predict potential risk areas for CCHFV across the Old World, with a special focus on Europe. Our model improved the prediction and attempted to give a more reliable and detailed map of habitat suitability for *H. marginatum*. As observed in the previous ecological niche model [63], the highest environmental suitability for *H. marginatum* was predicted across Southern Europe. In Southeastern part of Europe, the highest suitability was predicted across all Balkan countries. In contrast to the previous model [63], our prediction shows broader high and very high suitable areas particularly in Balkan countries (Albania, Croatia, Bosnia and Herzegovina, Kosovo, Montenegro, Romania, and Slovenia), Western Europe, United Kingdom and southern parts of Scandinavian countries which were underestimated by the study of Okely and colleagues [63], and less suitable areas in Central Europe. Besides the prediction of Okely and colleagues [63], previous studies have mapped the geographic distribution of *H. marginatum* based on regional and global scales [64,65]. The prediction of *H. marginatum* in our model suggested higher suitability for the occurrence of the species in Central, Western, Eastern and Northern Europe than the previous studies [64,65] which predicted low suitability for this tick species in this same region.

*H. marginatum* was first reported in Central Europe in Germany in 2007 [17]. Although *H. marginatum* was recorded in other countries of Central Europe, for example, the species was previously reported in Hungary [18], Slovakia [19], Czech Republic [19,21], and Austria [20]. These observations did not represent stable populations of *H. marginatum* in Central Europe. *H. marginatum* ticks have ecological plasticity that can support a wide range of tolerance to temperature and humidity conditions [66]. *H. marginatum* relatively preferred dry and warm regions [67], so, all stages of *H. marginatum* are most abundant during the summer months. *H. marginatum* is activated in the northern hemisphere when the temperature increases in spring, usually at the beginning of April [68]. Immature stages of *H. marginatum* are active between June and October, with the highest numbers in July and August. Following a blood meal, immature ticks either drop in early summer, molt to adults during the same season and overwinter as an adult or drop in late summer and overwinter as nymphs, molting to adults the following spring [68].

Stable populations of *H. marginatum* in Europe are restricted to the warm areas of the Mediterranean basin and are absent in Central Europe, likely due to environmental conditions. The ENM of *H. marginatum* anticipated its distribution in all countries across Central Europe. Our model provided strong evidence for occurrence of extensive medium to very high suitability in Germany, Poland, Hungary, and Czech Republic followed by Austria and Slovakia. Recently, the study that assessed seroprevalence of CCHF in Hungary showed seropositivity and Hungary can be considered as a potentially new geographical area in the distribution of CCHFV [69]. Moreover, some sporadic occurrence records of *H. marginatum* in other countries of Central Europe aside from Hungary shows that there can be potentially suitable habitats for breeding of *H. marginatum*. In Austria, the first *H. marginatum* (adult male extracted from a horse) occurrence was recorded in 2018 in Melk district in the state of Lower Austria [20]. Recent studies in Czech Republic and Slovakia reported *H. marginatum* records only on migratory birds [19]. Besides, very recent study reported five adult *H. marginatum* collected from horses and household in Czech Republic [21]. Although *H. marginatum* do not belong to endemic tick fauna in Germany, it has been found sporadically in recent years (e.g., 2007, 2011, 2017, and 2018) [70]. In Poland, *H. marginatum* records dated back to earlier times where one specimen of unfed *H. marginatum* male was reported in 1935 and three specimens in 1943 in Bytom, Upper Silesia, which are archived in Bytom’s Museum collection. Only two records from migratory birds have been noted to date: one on *Motacilla flava* [71] and one on *Acrocephalus schoenobaenus* [72]. It is very likely that these adult ticks were introduced to the Central Europe as nymphs feeding on migratory birds and moulted to adults after full engorgement. It appears that the environmental conditions in Central Europe provide the suitable habitat for *Hyalomma* spp. to complete their development and to find suitable vertebrate hosts for subsequent blood feeding. It has been suggested that *H. rufipes*, another important vector of CCHFV, overwinters in Central Europe [73], however, there is no speculation for establishment of *H. marginatum* populations.

Even though *H. marginatum* ticks have relatively low mobility by themselves, they can be transported over great distances by their vertebrate hosts, particularly migratory birds, and ungulates. Migratory birds have a key role in transportation of immature stages of *H. marginatum* ticks and can lead to the northward spread of *H. marginatum* and the establishment of permanent populations. During the spring migration of migratory birds from the south to the north, *Hyalomma* ticks are introduced to Central Europe. The spread of these ticks into new geographic locations can also cause the emergence of CCHFV. The presence of the virus, its vectors, reservoirs, and amplifying hosts are necessary for the emergence of CCHF, but suitable environmental conditions are also essential [28].

Appropriate climatic and biotic conditions in the regions that our map depicted may provide a suitable environment for introduced *H. marginatum* ticks. Particularly, a trend towards a warmer climate may be more favourable for maintaining infected *H. marginatum* ticks. Based on climate scenarios developed for the countries in central part of Europe, an expansion of climatically suitable habitats is expected in the near future for *H. marginatum*. Thus, it can be speculated that climate change will unquestionably increase the winter temperatures, leading to the increase of the probability for overwintering of *H. marginatum*, and consequently increasing the risk of establishment of *H. marginatum* in several parts of the countries in Central Europe. Therefore, modeling studies based on future scenarios should be implemented to better understand climate change impacts on *H. marginatum* dispersion.

Furthermore, our study provides evidence for strict-extrapolation risks between calibration and projection areas via the MOP metric. It is crucial to identify areas with high associated uncertainty in model projections to avoid misinterpretations of the potential distributions of disease vectors and to better interpret the prediction maps [49]. The MOP analysis is, therefore, a useful tool for dealing with these problems by performing robust identifications of extrapolation risks [51]. MOP results indicated broad areas of strict extrapolation in Southwest and Northwest China, and only small areas of strict extrapolation were detected in East Africa. Since these areas showed high to moderate model uncertainty in the analysis, the distribution prediction of *H. marginatum* in these areas should be taken with caution (**Figure 5**).

ENM results are useful in understanding species tolerances to biotic and abiotic factors, which, together with knowledge about ecology, behavior, and life history, aid in the selection of the most realistic predictions [74]. Our prediction for current global potential distribution of this tick species can help in understanding disease risk areas associated with this vector. However, assessing uncertainty in model projection is critical for vector and disease surveillance researchers and public health decision-makers. In summary, our future studies will consider further detailed mapping of CCHF disease and *H. marginatum* distribution under different climate change scenarios future projections.

## Acknowledgements

The authors would like to thank all resources and support provided by the Ain Shams University Disease Ecology and Dynamic Laboratory and the Ain Shams University Department of Entomology.

## Notes

### Competing Interest Statement

The authors have declared no competing interest.

